# ETV5 expression positively correlates with promoter methylation and predicts response for 5-FU-based adjuvant therapy response in proximal colon cancer

**DOI:** 10.1101/2020.09.08.287953

**Authors:** Anil K Giri

## Abstract

Discovery of markers predictive for 5-Fluorouracil (5-FU)-based adjuvant chemotherapy (adjCTX) response in patients with locally advanced stage II and III colorectal cancer (CRC) is necessary for early identification of potential responders as only 20-65% of CRC patients benefit from the treatment. PEA3 subfamily of ETS transcription factors (ETV1, ETV4, and ETV5) are upregulated in multiple cancers including colon cancers. However, the underlying epigenetic mechanism regulating their overexpression and their role in predicting therapy response in colon cancer is largely unexplored. In this study, using gene expression and methylation data from The Cancer Genome Atlas (TCGA) project, we showed that promoter DNA methylation negatively correlates with ETV4 expression (ρ= -0.17, p=5.6×10^−3^) and positively correlates with ETV5 expression (ρ= 0.22, p=1.43×10^−4^) in colon cancer tissue. Further, our analysis in 662 colon cancer patients treated with 5-FU-based-adjCTX revealed that higher ETV5 expression associated with shorter relapse-free survival (RFS) of treated patients with proximal tumors (Hazard ratio = 3.30 - 6.22, p=0.005-0.02). We also observed higher expression of signaling molecules involved in cellular proliferation (e.g. GNB5, DUSP4, FYN) in patients with high ETV5 level, suggesting that the increased cellular proliferation due to overexpression of these genes could drive the therapy resistance. The present study suggests ETV5 expression as a strong predictive biomarker for 5-FU-based adjCTX response in stage II/III CRC patients with proximal tumors.

## Introduction

Colon rectal adenocarcinoma (CRC) is the fourth most commonly diagnosed cancer (2 million cases in 2018) in the world and kills nearly 1 million people annually^1^ mostly due to the spread of tumor cells to other secondary organs (e.g. liver) in the later stage (stage III and IV) of the disease^2^. Therefore, successful treatment of the locally advanced early-stage (stage I, II, and III) cancer is necessary in order to prevent disease progression and improve the overall survival of the patients^3^. Usually, the early-stage patients are cured by surgical removal of the tumor only without the use of chemotherapy, however, systematic use of 5-Fluorouracil (5-FU)-based-adjCTX is recommended for stage II and III cases with high risk (e.g. with perineural invasion and poor histological differentiation) of reoccurrence^3,4^. However, the use of adjCTX after surgery cures only 20% of additional stage III patients over surgery alone (cures 50% of cases) and improves the chance of 10-year overall survival only by 10%–20% in stage II patients^4^. Further, it incurs considerable toxicity (e.g. myelosuppression, diarrhea) and economic costs to the patients^5, 6^. The higher toxicity and low efficacy of 5-FU-based adjCTX demand novel and reliable molecular markers that can predict the treatment response in early-stage (II and III) patients and help to stratify patients with different response^7,8^.

Attempts to predict response for adjuvant chemotherapy have identified molecular alterations (e.g. microsatellites status^9,^ TP53 mutations^10^, genetic polymorphism in MTHFR^11^ and DPYD^12,13^) as a predictive marker for 5-FU-based adjCTX response. Further, recent studies exploring gene expression signatures as predictive markers for treatment response in colon cancer have identified ESR1^14^, and CD8^15^ expression as predictors for 5-FU-based adjCTX response in CRC. However, none of the identified markers can successfully segregate the responders from nonresponders before initiating the therapy indicating the need to search for additional novel markers predictive for adjCTX response^16^.

Members of the polyoma enhancer activator 3 (PEA3) subfamily of E26 transformation-specific (ETS) domain-containing transcription factors which include ETV1, ETV4, and ETV5 promote cancer cell proliferation and survival in solid tumors including gastric ^17^, ovarian cancer^18^, and colon^19^ cancers and are being targeted for therapy^20^. However, the role of epigenetic mechanism especially DNA methylation in regulating their expression in colon cancer^18^ is largely unexplored. Further, the available evidence suggests that PEA3 subfamily members could be a potential biomarker for therapy response against CRC^18^, and their role in predicting adjCTX response has not been explored in any cancers including CRC. The current study explores the role of DNA methylation in regulating PEA3 members gene expression using The Cancer Genome Atlas (TCGA) colon adenocarcinoma cohort. Further, we explored the PEA3 subfamily ETS transcription factors as a predictive biomarker for 5-FU based adjCTX response in stage II and III CRC patients. Our analysis identified and validated ETV5 as a predictive marker for 5-FU-based adjCTX response in 662 stage II and II patients.

## Material and Method

### Processing of gene expression and DNA methylation data from the TCGA cohort

The expectation-maximization genes normalized RNA-Seq data for 328 colon adenocarcinoma patients samples (41 normal and 287 cancerous tissue) profiled in the TCGA project were downloaded using the Firehose tool (http://gdac.broadinstitute.org/). The data were further normalized using voom function in the limma package^21^, and Z-transformed before the differential and correlation analyses. We also downloaded methylation data for 482,481 CpGs using Infinium HumanMethylation450 Beadchip for 38 normal and 297 cancerous tissue samples. Methylation status at a CpG site was measured as beta value (β), which is the ratio of the methylated probe intensity and the overall intensity (sum of methylated and unmethylated probe intensities designed for a particular CpG in 450K beadchip). β ranges from 0 to 1, indicating no methylation (β = 0) to complete methylation of the CpGs (β = 1). In all the analyses, we performed appropriate quality control of the published data before their downstream analysis as described previously^7^,^22^. Briefly, we removed all the CpGs with missing values and CpGs assessed by probes that have a tendency of cross-hybridization, as specified in the supplementary file of Chen et al^23^. We used BMIQ normalization^24^ to remove any possible bias due to design differences in the type of probes (the type I and type II probes) present in the Illumina 450K platform before averaging of probes in the promoter region. The average methylation of probes between 1,500 bases upstream of the transcription start site (TSS) was defined as promoter methylation level for a gene.

### Discovery cohort

We downloaded the clinical and normalized gene expression profile of 477 stage II and III colon cancer patients collected under The French national Cartes d’Identité des Tumeurs (CIT) program from the Gene Expression Omnibus (GEO) platform (GSE39582)^25^. These primary patients with tumors have been treated with 5-Fluorouracil based adjuvant chemotherapy after surgery and monitored for relapse (distant and/or locoregional recurrence; median follow-up of 51.5 months). The recurrence-free survival (RFS) for the patients has been calculated as the time from surgery to the first recurrence. Patients have been staged according to the American Joint Committee on cancer tumor node metastasis (TNM) staging system^26^. The location of the tumor has been noted as distal and tumor based on the anatomical location. The gene expression data have been generated on Affymetrix U133 Plus 2.0 chips and normalized using the robust multi-array average method implemented in the R package affy. Gene expression was summarized as the average expression levels of all the probes of the genes and was used for differential and survival analysis.

### Validation cohort

For validation of the 5-FU-based adjCTX response prediction ability of ETV5, we downloaded the clinical and normalized gene expression data (GSE14333) for 185 CRC patients collected and published by Jorissen et al^27^ from the GEO platform. The gene expression data have been collected from specimens derived from primary carcinomas and snap-frozen in liquid nitrogen immediately after surgery for storage at −80°C. The patients have received standard adjuvant chemotherapy (either single-agent 5-fluorouracil/capecitabine or 5-fluorouracil and oxaliplatin) or postoperative concurrent chemoradiotherapy (50.4 Gy in 28 fractions with concurrent 5-fluorouracil) according to hospital protocols ^27^. The median follow-up for the patients was 37.2 months (range 0.9 to 118.6 months). Disease-free survival (DFS) was calculated as the duration from surgical operation to cancer recurrence, second cancer, or death from any cause. The grading for tumor stage has been determined using AJCC cancer staging manual^20^ and the position of the tumor has been noted as left, right, colon, and rectum. We categorized the cancers in the rectum and left colon as distal and in the right colon as proximal in origin.

### Differential expression analysis

5-FU-based-adjCTX treated patients with ETV5 expression > 3rd quartile was classified as ETV5-high group and with expression and < 1st quartile was classified as ETV5-low group. Differential expression analysis was performed using the Wilcoxon rank-sum test.

### Survival analysis

The effect of average ETV5 over RFS or DFS was determined using the Cox regression analysis. The hazard ratio has been calculated as the exponential of the regression coefficient obtained from the fitted regression model. The significance of the model was tested using the log-rank test.

### Enrichment analysis

The biological pathway enrichment for 22 genes against the human genome as the background was performed using Genecodis 4. 0.^28^ and FDR adjusted hypergeometric test p-values were used to identified enriched pathways.

### Statistical analysis

All the statistical analysis has been performed using R version 3.5.3. The Non-parametric Wilcoxon rank test has been used to compare the gene expression between two groups. The survival analysis has been performed using Cox-proportional regression as implemented in the “survival” package and the survival plots have been drawn using “ggplot” and “GGally” package in R.

## Results

### Cancer tissue has higher expression of ETV4 and ETV5 genes that correlate with promoter methylation in CRC patients

First, we compared the expression level of ETV1, ETV4, and ETV5 between the normal and cancerous tissue in TCGA data and observed higher expression of only ETV4 (Wilcox test P=4.90×10^−25^) and ETV5 (Wilcox test P=5.17×10^−9^) in tumor suggesting their possible role in tumor biology (Figure 1A, and B). Further, we checked the role of promoter DNA methylation over expression of ETV4 and ETV5 in the cancer tissue as promoter hypermethylation can shutdown gene expression. Our analysis revealed that promoter DNA methylation nominally differs between cancer and normal tissue for both ETV4 and ETV5 (Figure 1C, and D). Further analysis revealed that promoter methylation negatively correlates with ETV4 expression (ρ= -0.17, p=5.6×10^−3^) whereas positively correlates with ETV5 expression (ρ= 0.22, p=1.43×10^−4^) in cancer tissue suggesting that DNA methylation play a strong role in regulating ETV5 and ETV4 expression in colon cancer tissue (Figure 1E, and F).

**Figure 1:**
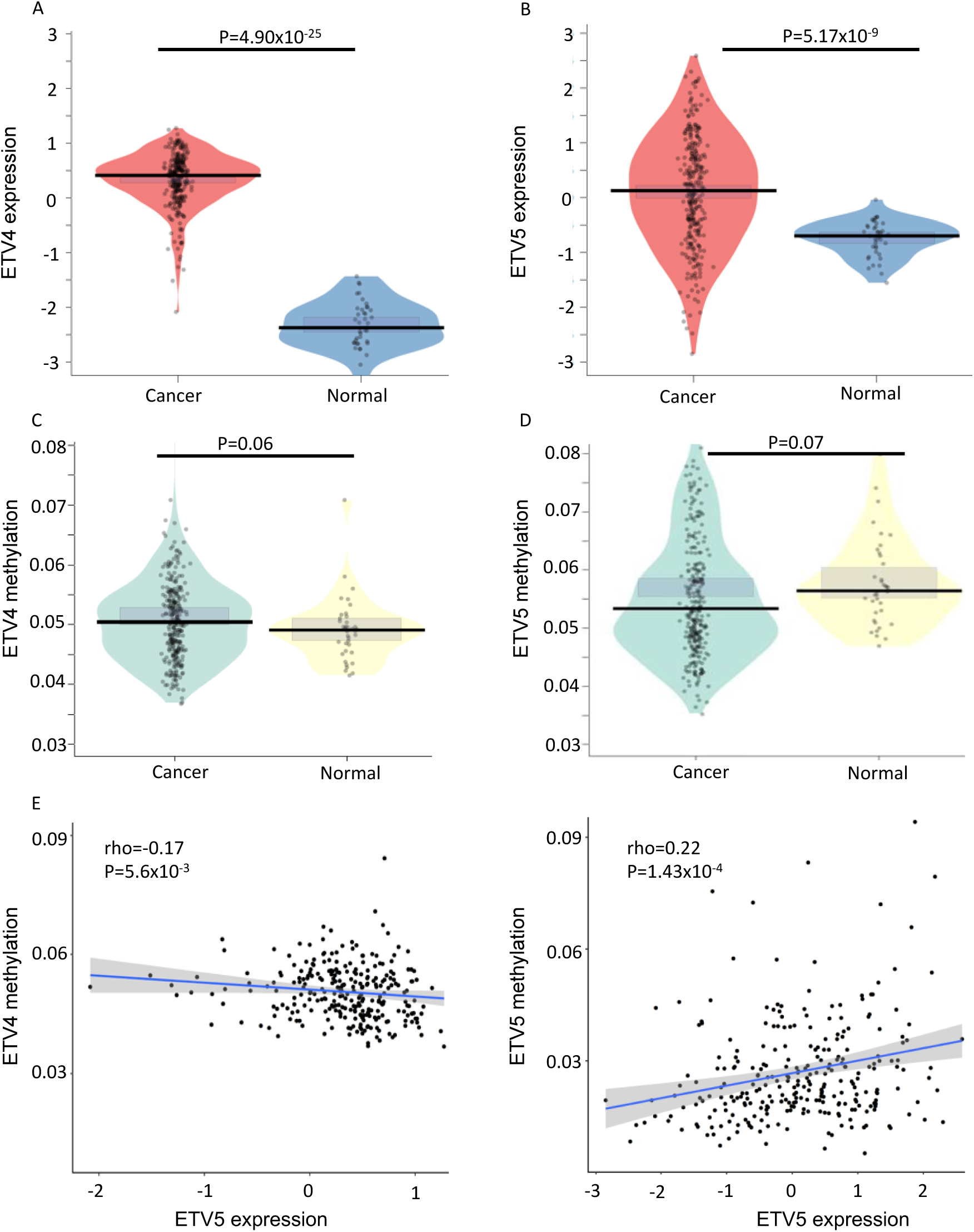
DNA methylation regulates ETV4 and ETV5 expression in colon cancer. ETV4 (A) and ETV5 (B) are overexpressed in cancerous tissue compared to healthy tissue in TCGA samples. Scatter plot showing promoter methylation difference between cancer and normal tissue for ETV4 (C) and ETV5 (D) genes. The p-value has been calculated using the Wilcoxon nonparametric test. Scatterplot showing spearman’s correlation between promoter methylation and expression levels of ETV4 (C) and ETV5 (D) in the cancerous tissue of CRC patients from TCGA data. The Spearman correlation coefficient and respective p-values have been shown in the figure.

### ETV5 expression differs in CRC patients with proximal and distal tumors

After observing that ETV4 and ETV5 are overexpressed in colon cancerous tissue from TCGA cohort, we studied their role in predicting adjCTX response in CRC using two publicly available datasets. Since the location of the tumor can affect the response to chemotherapy^29^, first we checked the expression level of ETV4 and ETV5 in stage II/III CRC patients (both treated and untreated) with proximal and distal tumors. We observed a significant difference in ETV4 expression between distal and proximal tumors only in discovery cohort samples (Figure 2A) but not in validation cohort (Figure 2B) whereas ETV5 expression differed both in discovery (Wilcox test P=8.05×10^−5^) and validation cohorts (Wilcox test P=0.005) (Figure 1B and C). The result suggests a tumor-side specific role of ETV5 in CRC. Further, in order to find the effect of ETV5 over disease progression, we compared the ETV5 expression level between adjCTX-treated patients with stage II and III and found no consistent significant difference across disease stages (Suppl. figure 1) in the discovery and validation cohorts (Suppl. Table 1).

**Figure 2:**
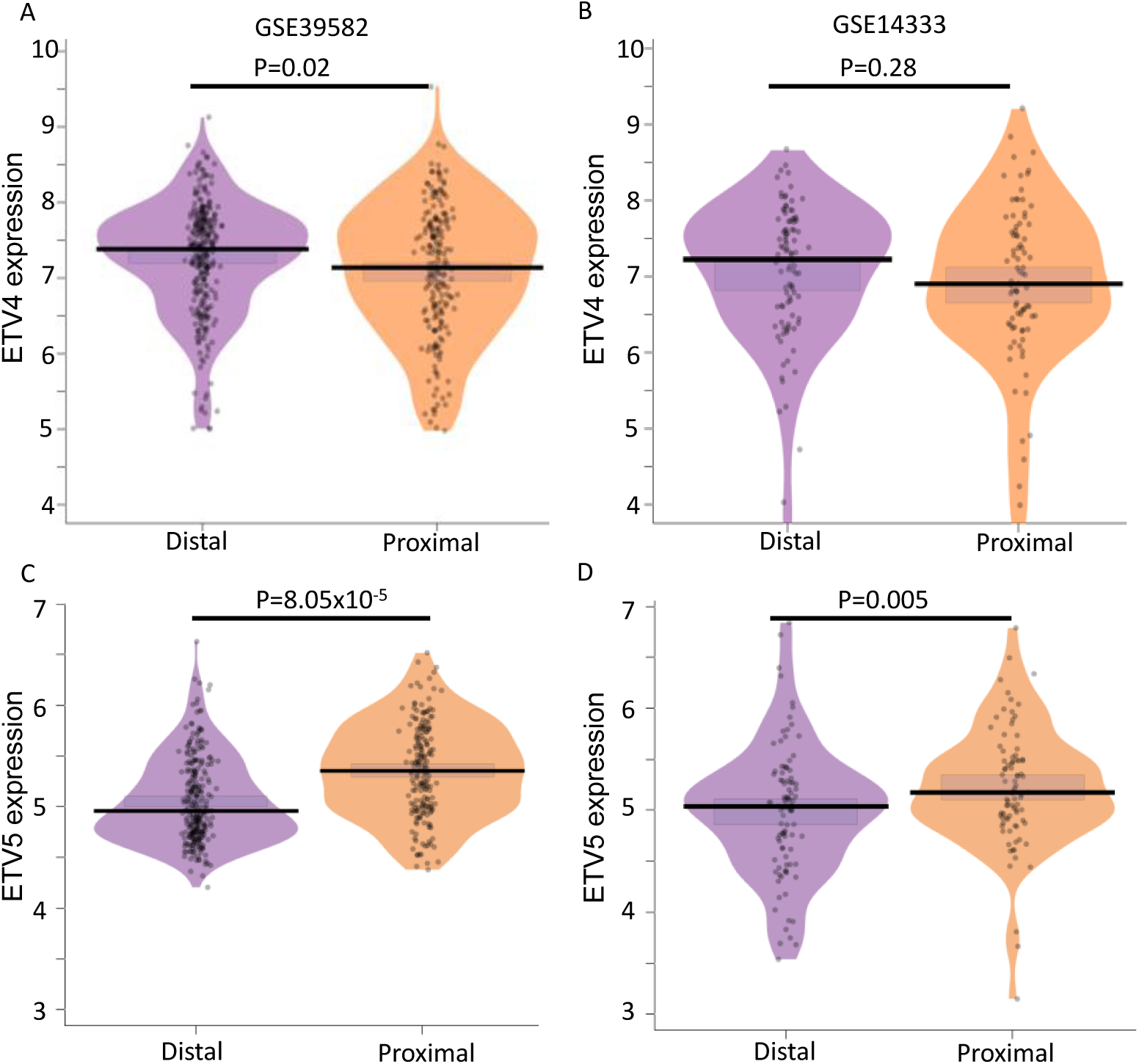
Comparison of ETV4 (A and B) and ETV5 (C and D) expression in distal and proximal tumors of combined stage II and III CRC patients from discovery (left side), and validation (right side) cohort. The P-value has been calculated using the Wilcoxon test.

### ETV5 correlates with disease free survival of stage II patients treated with adjuvant

In order to assess the effect of ETV4 and ETV5 expression over chemotherapy response, we used Cox-regression with RFS as the treatment outcome and observed that higher ETV5 expression is a strong and selective predictor for poor RFS (HR= 2.29, P=0.00178) in treated patients in the discovery cohort (Figure 3A). Further, stratified analysis revealed that ETV5 expression significantly predicts for RFS of patients with both distal (HR=2.21, P=0.01) and proximal tumors (HR= 3.30, p=0.02) in the adjCTX treated patients in the discovery cohort (Figure 3B and C). Furthermore, survival analysis in 85 stages II/III adjCTX treated patients from the validation cohort revealed that ETV5 expression significantly associates with poor RFS in patients with proximal tumors (HR=6.22, P=5.77×10^−3^) but not with distal tumor (HR=1.21, P=0.57) (Figure 3D, E, and F). We also checked the effect of tumor location (proximal or distal) over RFS of adjCTX-treated patients and did not observe a significant effect in both the discovery (HR=1.03, P=0.92) and validation (HR=0.68, P=0.06) datasets (Suppl. Fig. 2). The results suggest that the association of ETV5 with poor RFS in 5-FU based adjCTX treated patients with proximal tumors is not confounded by the survival difference due to the location of the tumor. We did not observe a significant association of ETV4 with RFS of patients either with distal or proximal tumors.

**Figure 3:**
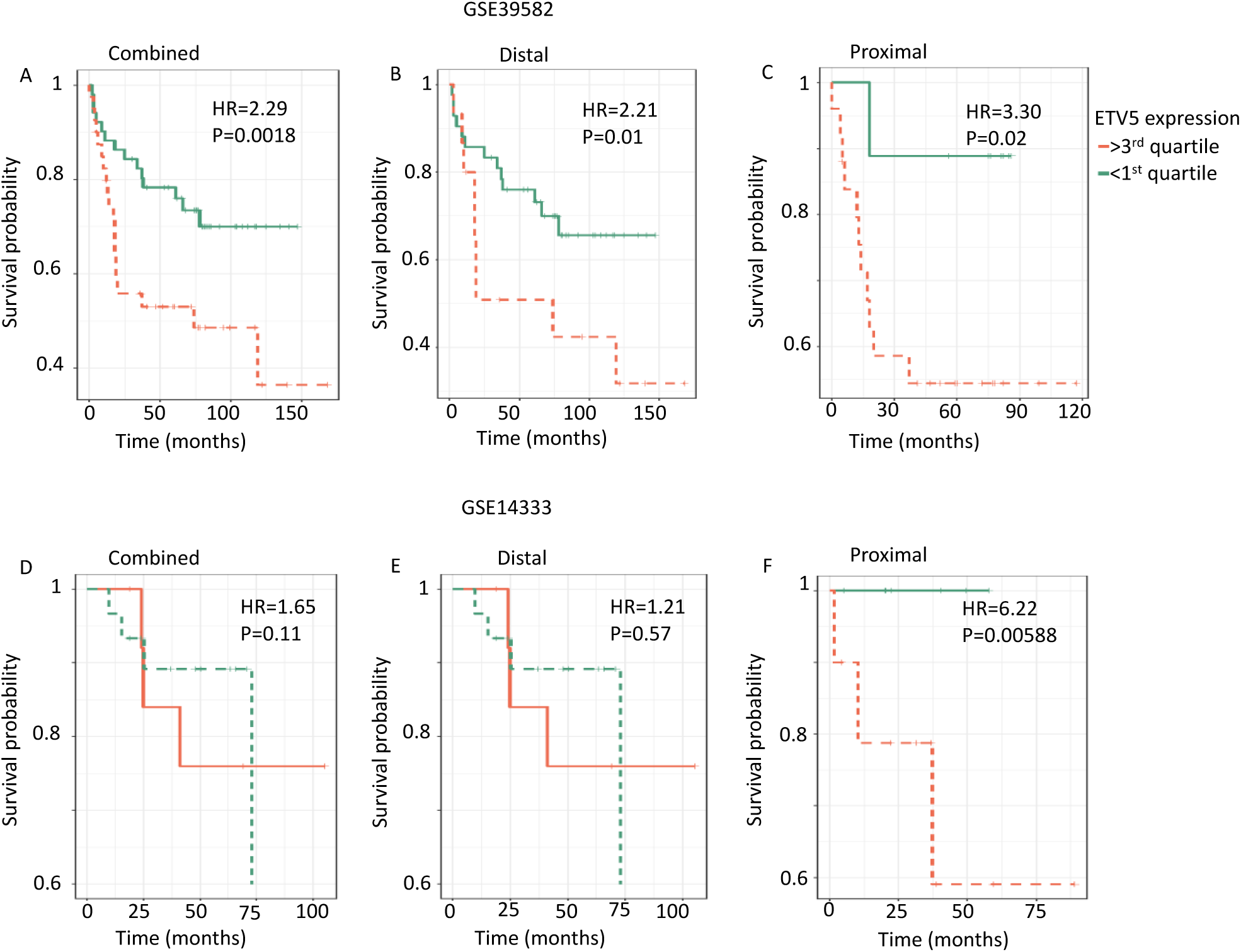
Effect of ETV5 expression over RFS in 5-FU-based adjCTX treated CRC patients. Kaplan– Meier survival curves for the probability of RFS of treated-CRC patients with low and high ETV5 has been shown for GSE39582-combined (A), distal, (B) proximal (C) tumors. Similarly, the probability for RFS of CRC patients in GSE14333 cohort has been shown for combined (D), distal (E), and proximal (F) tumors. The hazard ratio and p-value have been calculated using cox-regression and has been shown in the figure. For visualization purpose, patients with the ETV5 expression > 3^rd^ quartile has been categorized as high (red) and with expression <1^st^ quartile has been categorized as low (low). The log-rank analysis was used to test for significance between the survival curves.

### Higher ETV5 level associated with genes involved in cell differentiation

After observing that higher ETV5 expression correlates with shorter RFS in 5-FU-based adjCTX treated patients with proximal tumors. We explored the possible mechanism by which ETV5 may induce poor response for therapy. A differential analysis between adjCTX treated patients with higher (the 4^th^ quartile) and lower (the 1^st^ quartile) ETV5 expression revealed a strong difference (P < 10^−5^) in the expression of 22 genes enriched in the cell differentiation process (Suppl. Table 2) in both cohorts. The genes included signaling molecules (GNB5^30^, DUSP4^31^, FYN^32^), and drug transporter (ABCA3^33^) and 5-fluorouracil target genes (e.g. ACE2^34^) that have been already implicated in the resistance of 5-fluorouracil and other chemotherapies (Figure 4). The result suggests that ETV5 may induce drug resistance by activating signaling molecules and transcription factors involved in cell proliferation.

**Figure 4:**
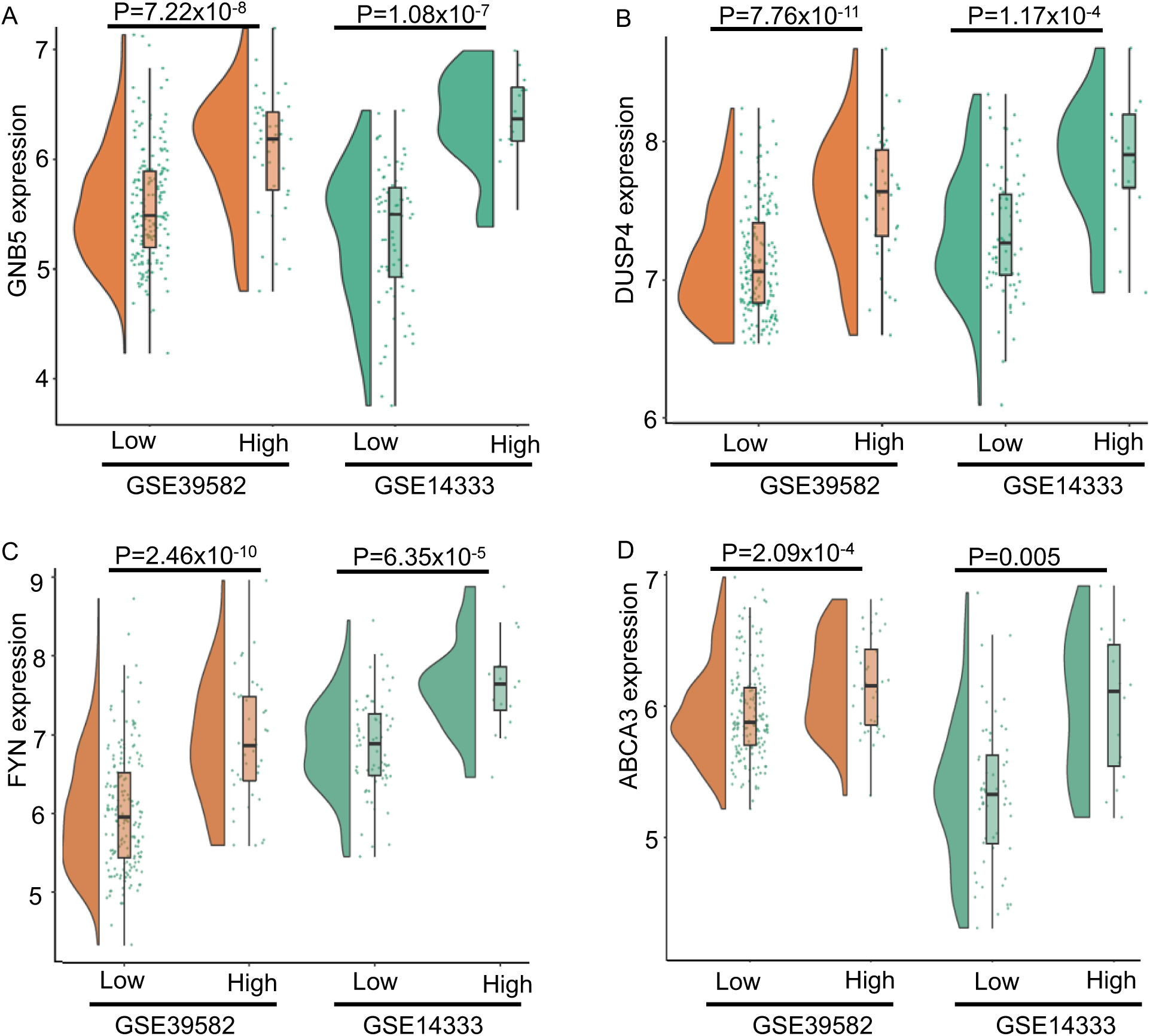
Higher ETV5 expression may induce therapy-resistance by higher expression of cell differentiation-associated genes. GNB5 (A), DUSP4 (B), FYN(C) andABCA3 (D) expression level in patients with high (>3^rd^ quartile) and low (<1^rd^ quartile) ETV5 expression in GSE39582 (left) and GSE14333 (right). P-value has been calculated using Wilcox nonparametric test.

## Discussion

5-FU-based adjCTX after surgery is the primary choice of treatment in early-stage colon cancer due to survival advantage over surgery alone^3,35^. However, higher toxicity and inability to segregate responders from nonresponders using available markers is a notable challenge to the clinical success of therapy. Therefore, there is an urgent need to identify additional predictive markers for the 5FU-based-adjCTX response in CRC. To address the need, we investigated the expression regulation of members of the PEA3 subfamily of ETS transcription factors by DNA methylation and the potential of their expression as a predictive marker for chemotherapy response in stage II and III colorectal cancer patients using 2 publicly available independent colon cancer datasets.

Our analysis revealed higher expression ETV4 and ETV5 genes in colon cancer tissue compared to normal tissue in TCGA samples (Figure 1A and B). Further, ETV4 expression showed a negative correlation with promoter methylation (Figure 1E) suggesting that DNA methylation mediates expression by hindering transcription factor binding^36^. Additionally, we observed a paradoxical positive correlation between ETV5 promoter hypermethylation and expression in cancer tissue suggesting that hypermethylation may facilitate gene expression either by the opening of chromatin^37^ or mechanical inhibition of transcriptional repressor binding^38^ or allowing transcription from an alternative promoter^39^. To our knowledge, there has been no study exploring the methylation-expression relation of PEA3 member proteins in colon cancers, and the results need to be validated in independent CRC cohorts. Further, a detailed mechanistic study in higher experimental model systems is needed on how the promoter hypermethylation increases the gene expression.

Additionally, we explored the role of ETV4 and ETV5 in 5-FU-based-adjCTX response prediction utilizing two publicly available CRC patient cohort. We observed higher expression of ETV5 in the proximal CRC tumor as compared to the distal tumor (Figure 2C and D) in both cohorts which is in accordance with more aggressive and high-grade histology of proximal tumors compared to distal tumors^40^ as higher ETV5 expression has been associated with faster cell proliferation and aggressive phenotypes^40^. Further, ETV5 overexpression stimulates CRC angiogenesis through activation of VGFR by PDGFR-β/Src/STAT3 signaling^18^ and increases bevacizumab resistance through AKT, ERK, and p38 signaling decreasing overall survival of CRC patients^18^. Survival analysis revealed that higher ETV5 expression significantly associated with shorter RFS in proximal colon cancer in stage II and III patients (Figure 3). Further, patients with higher ETV5 levels also had higher GNB5 expression (Figure 4A) which is a downstream Gbg subunit of G-protein coupled receptor (GPCR)^30^. GNB5 transduces the signal to various downstream effectors including RAS-MEK-ERK and PI3K-AKT pathways leading to cellular proliferation and viability^30^. GNB5 has emerged as the major target to overcome cetuximab resistance in colorectal cancer^30^. Furthermore, we observed an expression difference of DUSP4 in patients with a high level of ETV5. DUSP4 is a MAPK phosphatase and its role has been contradictory as downregulation of DUSP4 enhances cell proliferation and invasiveness in colorectal carcinomas^41^ but inhibits growth in human colorectal cancer cell lines^42^. These results suggest that higher ETV5 expression may induce drug resistance by upregulation of genes involved in colon cancer proliferation and growth. However, studies in larger human cohort and animal model systems can fully explain the detailed mechanism for the ETV5 role in 5-FU-based adjCTX resistance.

The current study identified ETV5 as a biomarker of 5-FU-based adjCTX response in CRC patients with evidence II level as defined by Simon et al^43^, and revealed that higher ETV5 is associated with poor response in proximal CRC tumors. These results suggest that ETV5 could be useful for the identification of responders before administration 5-FU-based adjCTX when included along with other already established clinicopathological markers.

## Author contribution

AKG conceptualized the study, analyzed the data, and wrote the manuscript.

## Acknowledgment

The author is very much thankful to Prof. Tero Aittokallio and Aleksandr Ianevski for their valuable suggestions during the analysis and editing of the manuscript.

---

**Suppl. Table 1:** Clinical characteristic of studied patients

|  | Discovery (GSE39582) |  | Validation (GSE14333) |  |
| --- | --- | --- | --- | --- |
| Traits | Treated (210) | Untreated (267) | Treated (85) | Untreated (100) |
| Age (<60/>60) | 109/101 | 64/203 | 38/47 | 19/81 |
| Sex (M/F) | 116/94 | 152/115 | 45/40 | 53/47 |
| Location (distal/proximal) | 137/73 | 144/123 | 46/39 | 54/46 |
| Stage (II/III) | 58/152 | 210/57 | 22/63 | 72/28 |

**Suppl. Table 2:** Biological process enrichment analysis for 22 common genes differentially expressed in ETV5 high vs low patients in both discovery and validation cohort. FDR corrected p-value for hypogeometric test for enrichment has been shown in the table.

| SN | Process | Hyp_pval_adj | Genes |
| --- | --- | --- | --- |
| 1 | Cell differentiation | 2.18x10<sup>-4</sup> | CDX2, SOX13, FYN, MECOM, ETV5, ETV1, DPYSL2 |
| 2 | Multicellular organism development | 2.11x10<sup>-3</sup> | CEBPA, CDX2, FYN, MECOM, DPYSL2, ANO1 |
| 3 | Negative regulation of transcription | 1.57x10<sup>-4</sup> | CEBPA, CDX2, RUNX1 |
| 4 | Response to drug | 6.02x10<sup>-4</sup> | FYN, ABCA3, DPYSL2 |

**Suppl. Fig. 1:**
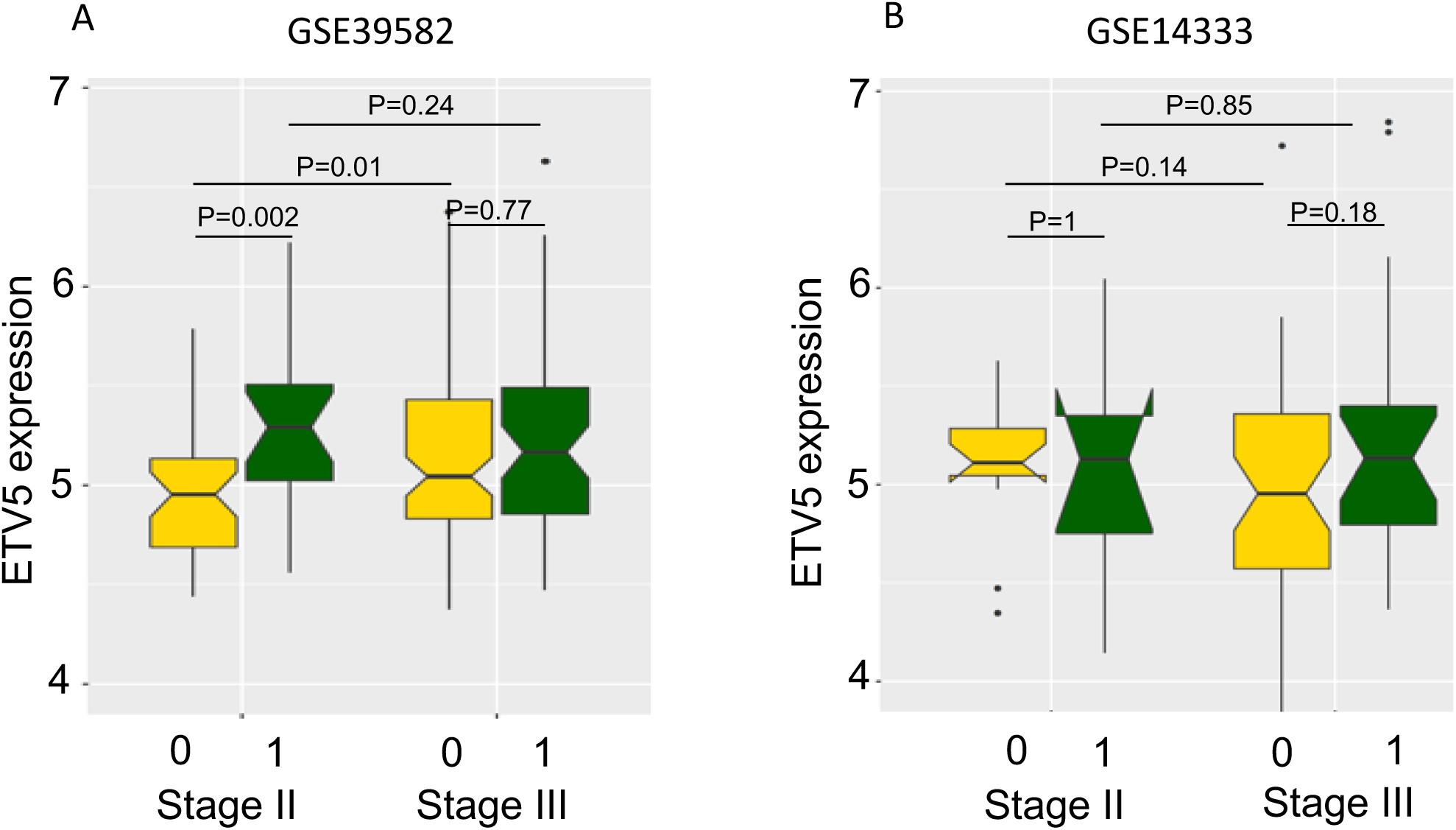
Comparison of ETV5 expression across different stages of relapse (coded as 1) and non-relapse (coded as 0) patients in adjCTX treated patients from GSE39582(A) and GSE1433 cohort (B). P-values have been calculated using Wilcox nonparametric test.

**Suppl. Fig. 2:**
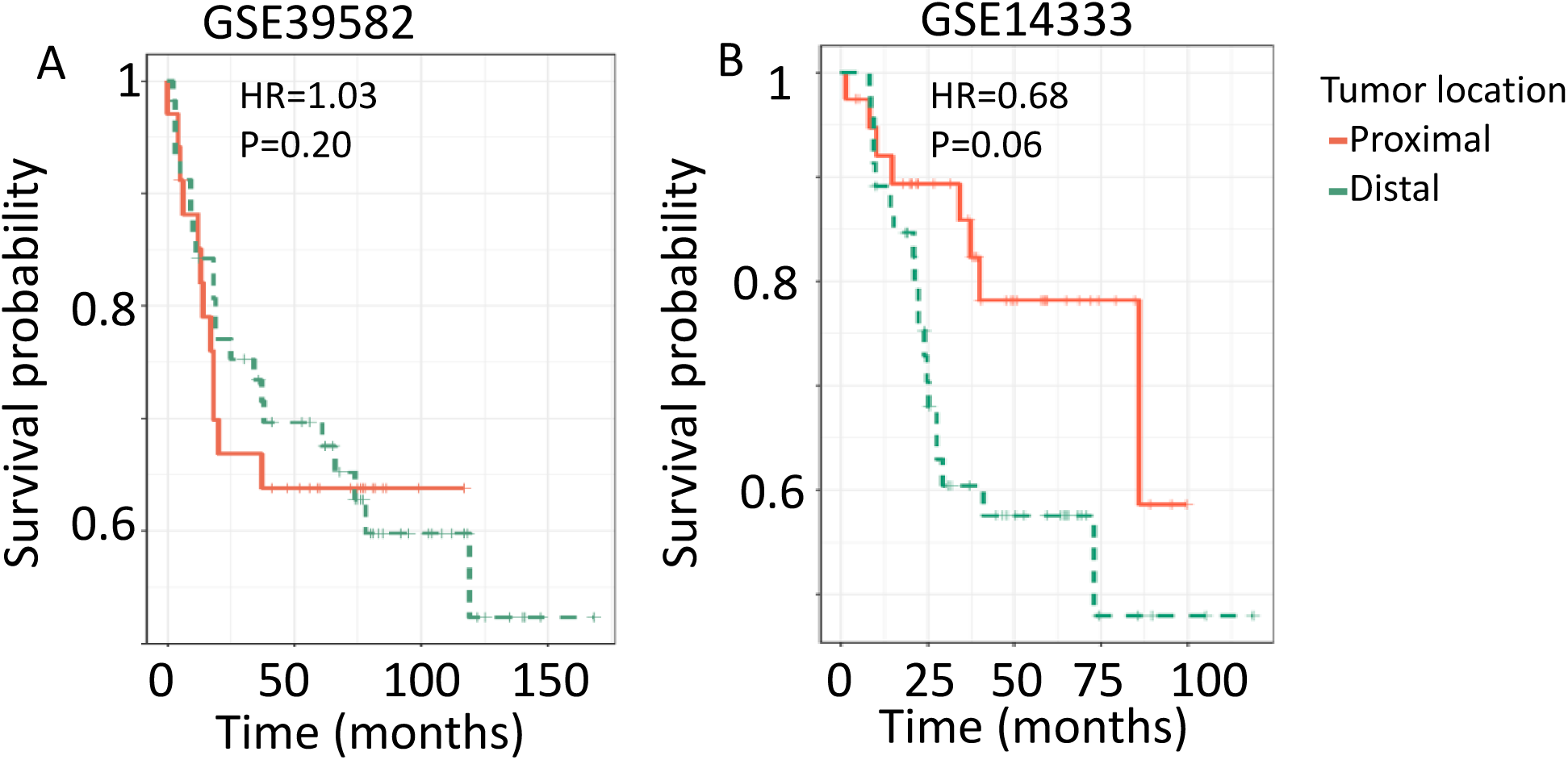
Effect of tumor location on the survival of the patients treated with 5-FU-based adjCTX in GSE39582 (A) and GSE14333 (B) cohort. The hazard ratio and p-value from the cox-regression has been shown in the figure.

